# Decoding Subject’s Own Name in the Primary Auditory Cortex

**DOI:** 10.1101/2022.07.30.502169

**Authors:** Hang Wu, Dong Wang, Yueyao Liu, Musi Xie, Liwei Zhou, Yiwen Wang, Jin Cao, Yujuan Huang, Mincong Qiu, Pengmin Qin

**Author notes:** Correspondence to: Dr. Pengmin Qin, Centre for Studies of Psychological Applications, Guangdong Key Laboratory of Mental Health and Cognitive Science, School of Psychology, South China Normal University, Guangzhou, Guangdong, 510631, China.

## Abstract

Current studies have shown that perception of subject’s own name (SON) involves multiple multimodal brain regions, while activities in unimodal sensory regions (i.e., primary auditory cortex) and their interaction with multimodal regions during the self-processing remain unclear. To answer this, we combined multivariate pattern analysis and dynamic causal modelling analysis to explore the regional activation pattern and inter-region effective connection during the perception of SON. We found that SON and other names could be decoded from the activation pattern in the primary auditory cortex. In addition, we found an excitatory effect of SON on connections from the anterior insula/inferior frontal gyrus to the primary auditory cortex, and to the temporal parietal junction. Our findings extended the current knowledge of self-processing by showing that primary auditory cortex could discriminate SON from other names. Furthermore, our findings highlighted the importance of influence of the insula on the primary auditory cortex during self-processing.

## 1. Introduction

In recent years, the neural basis of the self has been an important area of research in neuroscience (Frewen et al., 2020; Hu et al., 2016; Qin and Northoff, 2011; Qin et al., 2020). Using self-related and non-self-related stimuli, previous brain imaging studies have shown that extensive multimodal brain regions are involved in self-processing, including the temporal parietal junction (TPJ), the anterior insula (INS), the pregenual anterior cingulate cortex (pACC), the posterior cingulate cortex (PCC), and the anteromedial prefrontal cortex (AMPFC) (Qin et al., 2012; Tacikowski et al., 2011, 2013; Wuyun et al., 2014). The above findings indicated the importance of multimodal brain regions in the processing of self-related stimuli. While recent studies on other high-level processing (e.g., semantic processing) found extensive involvement of unimodal sensory regions (i.e., primary auditory cortex) (Fedorenko et al., 2012; Salmi et al., 2017), in studies on self-processing, such involvement remains unclear. One possible reason for that is the univariate approach adopted by most previous self-studies relied on averaged signals to compare the response difference between different experimental conditions, which ignored the response patterns of voxels in specific brain regions (Weaverdyck et al., 2020). To this end, recent studies adopted Multivariate Pattern Analysis (MVPA) to decode the subtle hemodynamic activation patterns of task-specific brain regions (Kubilius et al., 2015). Using MVPA, studies have shown that several high-level cognitive process (e.g., semantic processing, or music-induced emotions) induced activities in multimodal regions as well as unimodal sensory region (i.e., primary auditory cortex) (Fedorenko et al., 2012; Putkinen et al., 2021; Salmi et al., 2017). However, evidence in the relationship between self-processing and the activation pattern of the primary auditory cortex is still lacking. Whether hearing subject’s own name (SON) and other names could be decoded from the response patterns in the primary auditory cortex remains unclear.

As an external stimulus with high self-relatedness, SON has been widely used to investigate the neural mechanism underlying self-processing (Northoff et al., 2011). By comparing SON with a stranger’s name or a familiar name (e.g. a friend’s name), current evidence has shown that SON could reliably induce several multimodal regions, in which the activation of the anterior INS and TPJ were consistently observed (Frewen et al., 2020; Nakane et al., 2016; Tacikowski et al., 2011). The anterior INS and TPJ were found to respectively process internal and external environment information to the self (Eddy, 2016; Seth, 2013). According to a recent meta-analysis, the anterior INS in particular, was found consistently involved in all three levels of self-processing (including Interoceptive-processing, Exteroceptive-processing and Mental-self-processing), suggesting its crucial role for the self (Qin et al., 2020). Furthermore, previous studies have shown that the TPJ is associated with self-other distinction (Knyazev et al., 2021; Tan et al., 2022). However, studies to date have not answered the question of how the TPJ and anterior INS interact during the perception of SON. More importantly, if the primary auditory cortex does have a distinctive response to SON, it is unlikely an isolated process, but through the interaction with self-related multimodal regions (e.g., anterior INS and TPJ). Whether and how this interaction takes place is yet to be investigated.

The purpose of the current study is two-fold: 1) to investigate whether the primary auditory cortex has a self-specific response when hearing SON; 2) to explore the effective connectivity among the primary auditory cortex, anterior INS, and TPJ, during the perception of SON. For that, a total of 32 healthy subjects were recruited, who were presented with four auditory stimuli, including SON, a friend’s name (FN), an unknown name (UN), and a sound clip with inversed sound waves of the SON (SONREV). Firstly, a univariate analysis was performed using General Linear Model (GLM), to validate whether anterior INS, TPJ and other regions were activated by SON as compared with the other three conditions. Secondly, to identify self-specific regions activated by SON, a whole-brain four-class searchlight analysis was firstly performed, where identified brain regions were then taken as ROIs for further ROI-level MVPA analysis. Finally, combing the results from the univariate analysis and MVPA, a dynamic causal modeling (DCM) analysis was conducted to investigate the modulatory effects of SON on the effective connectivity among the multimodal (i.e., anterior INS, TPJ or other regions revealed by univariate analysis) and unimodal sensory regions (i.e., primary auditory cortex or other regions revealed by MVPA). By doing so, we expected to combine the information from activation strength (i.e., GLM) and activation pattern (i.e., MVPA), to provide new evidence about how the primary auditory cortex responds to SON and how the functional dynamics among unimodal sensory region and multimodal regions were modulated by the SON.

## 2. Materials and Methods

### 2.1 Subjects

Thirty-two right-handed adults were recruited in this study (male/female: 15/17; age range: 18-25 years). Four subjects were excluded due to excessive head motion during the scanning (i.e. over 3 mm in translation or 3 degrees in rotation), leaving 28 subjects (male/female: 13/15; age range: 18-25 years) in the final analysis. All subjects had normal hearing, and reported no history of neurological or psychiatric disorders. Informed consent was obtained from each subject and ethical approval was obtained from the Research Ethics Board of the School of Psychology in South China Normal University.

### 2.2 Stimuli

Four types of auditory stimuli were used, including SON, FN, UN, and SONREV. In addition, to maintain the subjects’ attention, an English name “Jack” was also presented. All names were disyllables. For the FN, to minimize the confounding effects between self-relatedness and familiarity (e.g. in a close friend’s name or a parent’s name), the name of classmates provided by the subjects were used (Qin et al., 2012). Each subject’s UN was randomly selected from the other subjects’ names, to control the physical features of SON and UN at a group level. Furthermore, the SONREV was presented for each subject as a control condition, which has the same acoustic complexities and voice identities as the SON. In the current study, only FN and UN with the same gender of the subjects were chosen.

All voice recordings were edited using a sample rate of 44 100 Hz and a resolution of 16 bits, which were normalized by peak amplitudes (Zhang et al., 2022). The names used in this study were delivered by a native Chinese speaker, an adult female unknown to the subjects. The names were presented binaurally using headphones at 100 dB with E-prime 3.0 (Psychology Software Tools, Inc., Sharpsburg, Pennsylvania, USA), with a mean duration of 508 ± 52ms (SD). During scanning, all subjects confirmed that they could clearly hear the names.

### 2.3 Experimental Design

In a block design, subjects were presented with the four auditory stimuli (SON/FN/UN/SONREV). For each subject, 12 runs of functional image data were collected. For each run, two blocks of each of the four stimuli were presented (eight blocks in total). Each 12 s block comprised of 6 stimulus repetitions, with a 16 s inter-block interval. Furthermore, an English name (Jack) was presented alternately to maintain the subjects’ attention. During the experiment, subjects were instructed to keep their eyes open and concentrate on a gray cross presented on a black screen, while listening to the names passively and pressed a button with their right index finger when they heard “Jack”.

### 2.4 fMRI data acquisition and preprocessing

MR images were acquired on a 3T Siemens MAGNETOM Prisma-fit scanner (Erlangen, Germany). A high-resolution T1-weighted anatomical image (TR/TE/θ = 2530ms/2.3ms/7°, FOV = 256mm, matrix size = 256 × 256, slice thickness = 1 mm, 208 slices) was firstly acquired for each subject for image registration and localization. Functional images were then acquired using a T2*-weighted EPI sequence (TR/TE/θ = 2000ms/30ms/90°, FOV = 192mm, matrix size = 64 × 64, slice thickness = 3 mm, 32 slices).

Functional data were preprocessed using the afni_proc.py program in the AFNI software (version 20.1.07), which includes: slice timing correction; head motion correction using realignment on functional volumes; co-registration of high-resolution structural image with functional images; non-linear transformation to Montreal Neurological Institute (MNI) template; functional volumes resampled to 3 × 3 × 3mm^3^; volumes with derivative values of a Euclidean Norm (square root of the sum squares) above 0.3 in their six-dimensional motion parameters were censored in the following analysis, along with its previous volume; spatial smoothing was applied using a Gaussian filter with a full width at half maximum of 6 mm; and functional time series were scaled to percent signal change.

### 2.5 Univariate analysis

After preprocessing, univariate analysis was performed by the SPM12 (version 7771) scripts implemented in MATLAB 9.6 (R2019a; MathWorks). Firstly, a voxel-wise first-level analysis for each subject was adopted using a general linear model (GLM) with condition onsets and durations. The design matrix consisted of 11 regressors, including 4 experimental conditions (SON, FN, UN, SONREV), 6 motion parameters (3 rotations, 3 translations), and 1 binarized vector for censored volumes (a vector where 0 indicated censored time points, and 1 indicated non-censored time points). All regressors were produced by convolving canonical hemodynamic response function (HRF). The first-level results from all subjects were then used in a second-level (group level) analysis, in which a whole-brain one-way repeated measure ANOVA was firstly performed to analyze the main effect among the four conditions. The resulting whole-brain statistical maps were then controlled at a cluster-level significance threshold of p < 0.05 after FWE correction (p < 0.005 uncorrected, cluster size > 59 voxels). Based on the corresponding brain regions in the above ANOVA analysis, post-hoc ROI-based analyses were then performed to further test the difference between each pair of the four conditions. Specifically, for each ROI, a mean estimated coefficient was obtained in each condition respectively for each subject, which was then compared against the other conditions (SON vs. FN, SON vs. UN, SON vs. SONREV, FN vs. UN, FN vs. SONREV, and UN vs. SONREV), using two tailed paired-sample t-tests (Bonferroni corrected for multiple comparisons).

### 2.6 Multivariate pattern analysis

For each subject, a GLM was re-performed on the preprocessed (but unsmoothed) functional image, resulting in a total of 24 beta maps (2 block × 12 runs) per condition. The resulting individual beta maps were masked using a grey matter mask (i.e., non-cortical voxels were ignored) obtained from the AAL90 atlas, and normalized to zero mean and scaled to unit variance. Then a whole-brain four-class searchlight analysis was separately performed on these beta maps of each subject using a linear SVM classifier (implemented in the Nilearn package), with a sphere radius of 6mm. Since this is a multiclass classification problem (with four conditions, SON, FN, UN, and SONREV) which the linear SVM does not support, a one-vs-one (OVO) reduction scheme was used to transform it into binary classification, where each condition was discriminated from each of the remaining three conditions in a pairwise manner (Wurm and Caramazza, 2019). In addition, for cross validation, a leave-one-run-out method was conducted, where the classifier was trained on 11 runs and tested on the remaining one (Todd et al., 2013). This procedure was iterated 12 times, of which the resulting classification accuracy were averaged and assigned to the central voxel of each searchlight sphere, yielding an accuracy map (chance level at 25%) for each subject. These accuracy maps were then spatially smoothed using a Gaussian filter with 6 mm FWHM (Bulthé et al., 2014). To identify clusters with above-chance-level classification accuracies (25%), a non-parametric permutation test (analogous to a one sample t-test) was performed using the Randomise program in the FSL package, with 5000 permutations (Salmi et al., 2017) and a threshold of p < 0.05 (FWE corrected) for multiple comparisons.

To further investigate the brain regions that showed differential activation pattern especially between SON and non-SON (FN, UN, SONREV), an ROI-level MVPA analysis was performed based on the overlapping regions between the above searchlight result and AAL 90 atlas. For each ROI, six classification tasks (SON vs. FN, SON vs. UN, SON vs. SONREV, FN vs. UN, FN vs. SONREV, and UN vs. SONREV) were performed using a linear SVM classifier with a leave-one-run-out cross-validation, to obtain the average accuracy for each subject, which was then compared with the chance level (50%) using a one sample t-test, and controlled for multiple comparisons with the Bonferroni correction (p < 0.05).

### 2.7 DCM analysis

To investigate the effective connectivity among brain regions involved in the self-processing, and how it was modulated by SON, we used DCM 12.5 implemented in SPM12 (version 7771) on our fMRI data. Combining results from previous fMRI studies on self-processing (Qin et al., 2012; Tacikowski et al., 2011, 2013), a meta-analysis of the three-level model of self-processing (Qin et al., 2020), and our univariate and MVPA results, three brain regions were identified as ROIs in the DCM analysis: the primary auditory cortex, the anterior INS/inferior frontal gyrus (IFG), and the TPJ. The INS/IFG and TPJ were chosen due to a significantly higher activation in the SON than in the other three conditions in our univariate results, which is in line with previous studies (Qin et al., 2012; Tacikowski et al., 2013; Wuyun et al., 2014), while the primary auditory cortex, which was seldom focused in previous SON studies, was chosen due to its differential activation pattern between SON and non-SON stimuli in our MVPA results. For better interpretability, we built models for the left and right hemispheres respectively, and given that the right TPJ did not show significant activation in our univariate result, only the left hemisphere model was used in the following DCM analysis and reported in our main results. Results of the right hemisphere model were reported in the Supplementary Materials for validation.

As suggested by previous studies (Zhang and Du, 2022; Zhao et al., 2021), to minimize individual variations, we first created a 10 mm sphere (i.e., outer sphere) centered on the group peak coordinates, and then defined individual ROIs as 6 mm spheres (i.e., inner spheres) at the individual peak coordinate within the outer sphere. To reduce noise, only the significant voxels in the main effect of all four conditions were selected, using a threshold of p < 0.05 uncorrected. Finally, for each ROI, the first principal component of the BOLD time series was computed and adjusted by regressing out effects of no interest.

As suggested by previous studies (Putkinen et al., 2021; Willinger et al., 2021), a full connectivity model (‘full’ model) was specified for each subject for DCM analysis, which included three sets of parameters: effective intrinsic connections between the three regions and their self-connections (matrix A); modulatory effects of SON on all connections between regions and self-connections (matrix B); and driving inputs (using all four auditory stimuli: SON, FN, UN, SONREV) into the left primary auditory cortex (matrix C) (Zeidman et al., 2019a). Specifically, the self-connection of Matrices A and B reflected the self-inhibition in each region, i.e., their gain or sensitivity to driving inputs (Zeidman et al., 2019a). A positive value indicated an increase of regional self-inhibition due to the experimental manipulation, and a negative value indicated a disinhibition (Zhang and Du, 2022). To systematically investigate how the connections among these ROIs were modulated by the SON, all between-region and within-region connections for matrices A and B were switched on (Zeidman et al., 2019b). Furthermore, the driving inputs were not mean-centered, in which the A matrix represents the average connectivity estimated from unmodelled implicit baseline (i.e., the average connectivity of the FN, UN and SONREV), and the B matrix represents the modulatory effects of SON compared to the baseline activity (Bencivenga et al., 2021; Zeidman et al., 2019a).

The ‘full’ DCM model constructed was first used to perform the first-level (within-subject) analysis, in which the model was estimated for each subject using Bayesian model inversion, to find the best parameters (i.e., DCM model connections) that show a trade-off between accuracy (how closely the predicted timeseries correspond to the observed data) and complexity (how far the parameters have to move from their prior values to explain the data). Then group-level DCM inference was performed using the Parametric Empirical Bayes (PEB) framework (Zeidman et al., 2019b). By only inverting one DCM ‘full’ model for each subject, the PEB approach have the advantage of avoiding estimating multiple DCMs for the same subject, which could fall into different local optima (Zeidman et al., 2019b). Specifically, an automatic search procedure was adopted using the Bayesian model reduction (BMR) approach to iteratively prune any parameters that do not contribute to model evidence, leaving the parameters with the most evidence (Friston et al., 2016). Finally, Bayesian model Average (BMA) was used to calculate the average parameter values across the 256 models from the final iteration of the automatic search procedure, and a threshold based on the model evidence was used to keep only parameters with strong evidence, i.e., posterior probability > 0.95, which indicates the probability of a parameter being present vs. absent (Bencivenga et al., 2021; Zeidman et al., 2019b).

## 3. Results

### 3.1 Univariate analysis results

Firstly, in Figure 1A, significant main effect of all four conditions was found in five cortical regions, including bilateral INS/IFG, left TPJ, and bilateral primary auditory cortex (LAC and RAC) (p < 0.05, FWE corrected at cluster level with cluster size > 59 voxels). Secondly, in Figure 1B, further ROI-based post-hoc comparisons revealed that perception of the SON induced significantly stronger activation in bilateral anterior INS/IFG and the left TPJ compared with the other three conditions. In contrast, the SONREV induced a significant increase of activation in bilateral primary auditory cortex compared with the FN and UN, respectively. Bonferroni correction (p < 0.05) was used for multiple comparisons. Please see Table 1 for detailed results of the pairwise comparisons.

**Table 1.** Results of ROI-level Post-hoc Pairwise Comparisons

| Brain Region | T-value | Degrees of Freedom | P-value | Cohen's d |
| --- | --- | --- | --- | --- |
| LAC |  |  |  |  |
| SONREV vs. FN | 4.78 | 27 | < 0.01 | 0.38 |
| SONREV vs. UN | 4.50 | 27 | < 0.01 | 0.31 |
| RAC |  |  |  |  |
| SONREV vs. FN | 5.27 | 27 | < 0.01 | 0.57 |
| SONREV vs. UN | 4.48 | 27 | < 0.01 | 0.40 |
| LINS/IFG |  |  |  |  |
| SON vs. FN | 3.10 | 27 | 0.03 | 0.64 |
| SON vs. UN | 4.92 | 27 | < 0.01 | 1.11 |
| SON vs. SONREV | 4.05 | 27 | < 0.01 | 0.98 |
| RINS/IFG |  |  |  |  |
| SON vs. FN | 4.31 | 27 | < 0.01 | 0.76 |
| SON vs. UN | 3.91 | 27 | < 0.01 | 0.74 |
| SON vs. SONREV | 4.15 | 27 | < 0.01 | 0.88 |
| LTPJ |  |  |  |  |
| SON vs. FN | 3.84 | 27 | < 0.01 | 0.78 |
| SON vs. UN | 3.81 | 27 | < 0.01 | 0.59 |
| SON vs. SONREV | 3.74 | 27 | < 0.01 | 0.64 |

Note: AC = auditory cortex, INS = insula, IFG = inferior frontal gyrus, TPJ = temporal parietal junction.
SON = subject's own name, FN = familiar name, UN = unknown name, SONREV = a sound clip with
inversed sound waves of the SON. Bonferroni correction ( $p < 0.05$ ) was used for multiple comparisons.

**Figure 1.**
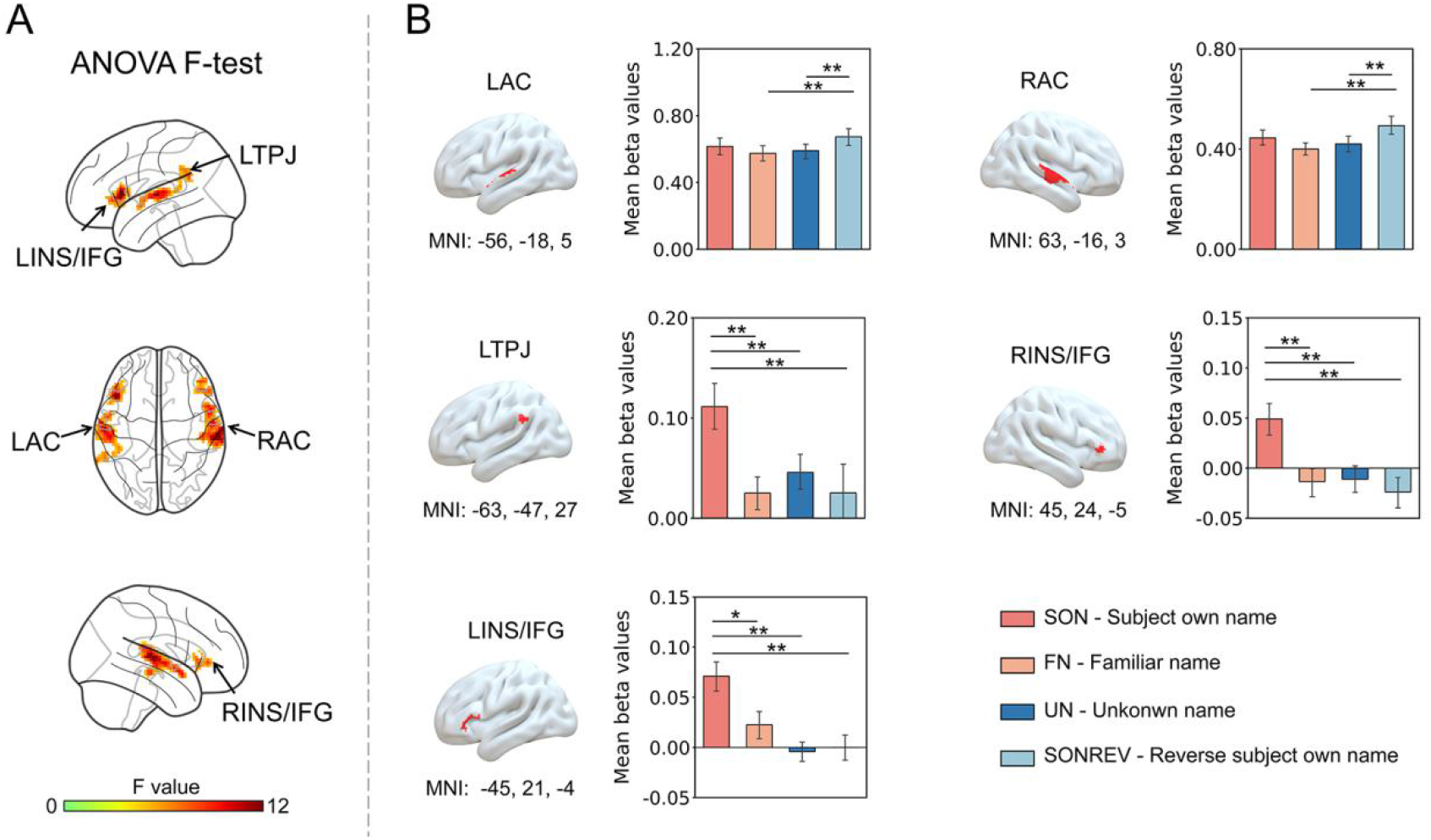
Brain regions with stronger activation for the SON compared with FN, UN, and SONREV. (A) Brain regions that showed a significant main effect among all four experimental conditions, resulting in five regions including bilateral primary AC (LAC and RAC), bilateral INS/IFG (LINS/IFG and RINS/IFG) and the left TPJ. (B) ROI-level post-hoc pairwise comparisons of the four experimental conditions were performed based on the above five brain regions, where SON consistently induced a higher activation in bilateral INS/IFG, and the left TPJ. AC = auditory cortex, INS = insula, IFG = inferior frontal gyrus, TPJ = temporal parietal junction. ** indicates p < 0.01 Bonferroni corrected; * indicates p < 0.05 Bonferroni corrected.

### 3.2 Multivariate pattern analysis results

For the whole-brain four-class searchlight analysis, the results showed a broad set of cortical areas with accuracy significantly above chance level (25%), including bilateral primary auditory cortex, the left IFG, and the left precentral gyrus (Figure 2A). Significance was calculated with a non-parametric permutation test and corrected for multiple comparisons using FWE correction (p < 0.05). To further examine whether the regions from the four-class searchlight analysis could consistently classify each pair of the four auditory conditions, especially the interested condition SON vs. non-SON (FN, UN, SONREV), respectively, an ROI-level MVPA analysis was performed based on the overlapping regions between the searchlight result and AAL 90 atlas. As seen in Figure 2B, the MVPA analysis revealed that bilateral primary auditory cortex showed a consistent above chance accuracy (50%) for classifying between SON and FN, SON and UN, as well as SON and SONREV (p < 0.05, Bonferroni corrected). For comparison results of the other three control conditions (ROI-level MVPA for classifying between FN and UN, FN and SONREV, as well as UN and SONREV), please see Supplementary Figure 1. Finally, in Figure 2C, a cortical map was shown to summarize the results from the univariate (cortical areas in blue) and ROI-level MVPA analysis (cortical areas in red), as well as their overlapping voxels (cortical areas in green), which identified three ROIs for further DCM analysis, including the left primary auditory cortex (from MVPA results), the left anterior INS/IFG (from univariate results), and the left TPJ (from univariate results).

**Figure 2.**
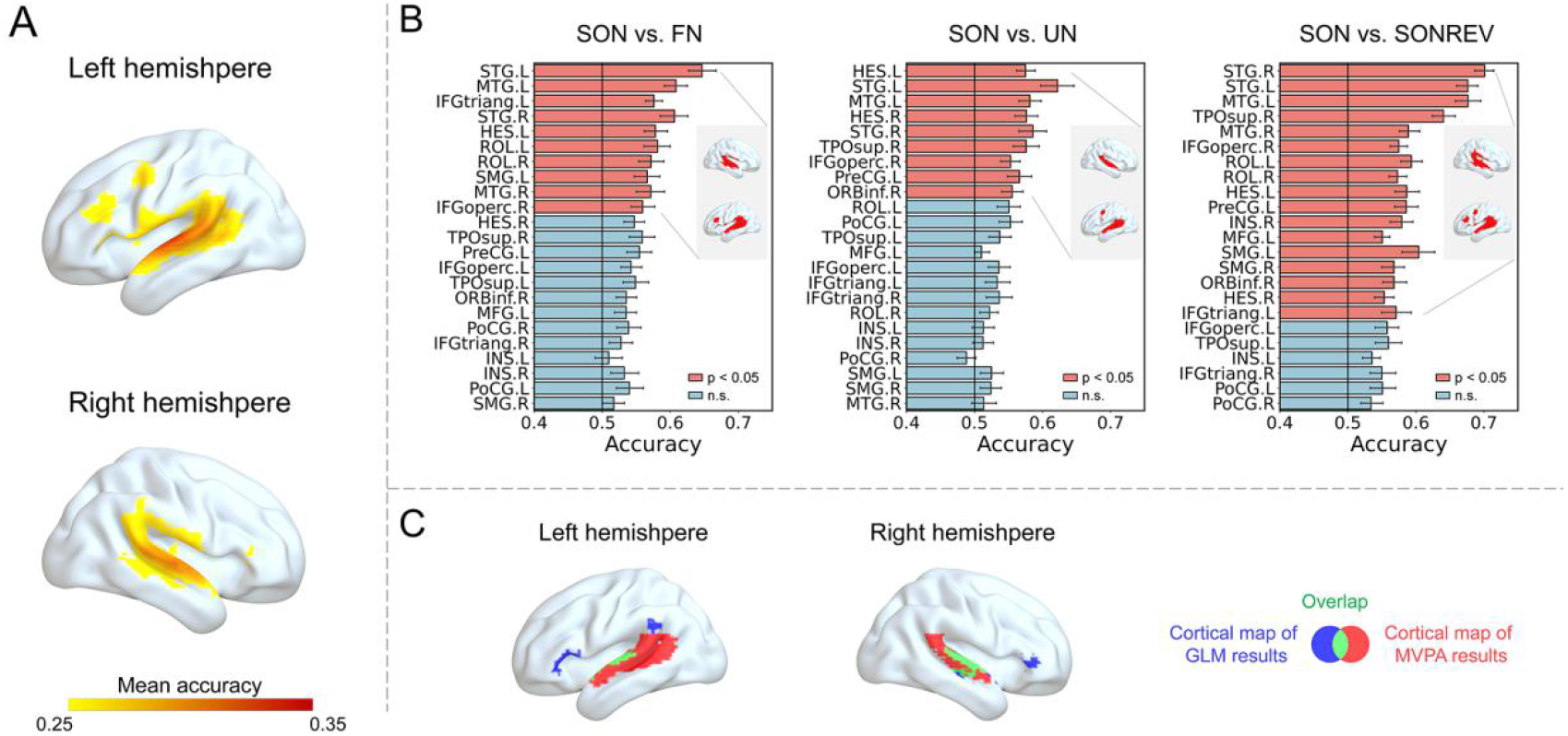
Brain regions that showed differential activation patterns between SON and non-SON conditions. (A) Results of four-class whole-brain searchlight analysis, in which voxels with classification accuracy above chance-level (25%) were displayed. (B) Based on the overlapping regions between the searchlight result and AAL 90 atlas, mean accuracy of ROI-level MVPA classification was presented for SON vs. FN, SON vs. UN, and SON vs. SONREV, respectively. For each classification, red bars indicate a mean accuracy significantly above chance-level (50%), and blue bars indicate the opposite. (C) Overlapped voxels (in green) between the results of activation strength (from univariate analysis, in blue) and activation pattern (from MVPA, in red). SON = subject’s own name, FN = familiar name, UN = unknown name, SONREV = a sound clip with inversed sound waves of the SON.

### 3.3 DCM analysis results

Based on the above analysis, DCM analysis was further conducted to explore the modulatory effect of SON on the effective connectivity among the left primary auditory cortex (MNI coordinates: -56, -18, 5), left anterior INS/IFG (MNI coordinates: -45, 21, -4), and left TPJ (MNI coordinates: -63, -47, 27). Firstly, a full connectivity model (‘full’ model) was constructed for each subject, where the three ROIs were bidirectionally connected with each other. Specifically, the four auditory conditions (SON, FN, UN, SONREV) were treated as driving inputs to the left primary auditory cortex, and SON was treated as the modulatory condition on all connections (Figure 3A). As shown in Figure 3B and Table 2, when compared with the other three conditions, SON enhanced the connectivity from the left anterior INS/IFG to the left primary auditory cortex (modulation: 1.08), and to the left TPJ (modulation: 0.43) with strong evidence (i.e., posterior probability > 0.95), which indicated an excitatory effect of the SON (Zeidman et al., 2019a). In addition, negative self-connection values were found in the left primary auditory cortex (modulation: -0.35) and left anterior INS/IFG (modulation: -0.98) during the perception of SON with strong evidence (i.e., posterior probability > 0.95), which indicated a disinhibition on these regions due to the SON (Zeidman et al., 2019a). For validation, we also performed DCM analysis of these three ROIs, in which SON was compared with FN, UN, and SONREV, respectively, and found consistent results as the main analysis (please see Supplementary Figure 2). In the Supplementary Materials, we also reported DCM results in the right hemisphere only, and found consistent results (please see Supplementary Figure 3 and Supplementary Table 1).

**Table 2.** Parameter Estimates of the Left Hemisphere

| Connection | Intrinsic connectivity | Posterior probability | Modulatory effect of SON | Posterior probability |
| --- | --- | --- | --- | --- |
| INS to TPJ | **0.16 (0.00)** | **1.00** | **0.43 (0.01)** | **1.00** |
| INS to auditory cortex | **0.38 (0.00)** | **1.00** | **1.08 (0.04)** | **1.00** |
| INS self-connection | **-0.49 (0.00)** | **1.00** | **-0.98 (0.01)** | **1.00** |
| TPJ to INS | 0.00 (0.00) | 0.00 | 0.00 (0.00) | 0.00 |
| TPJ to auditory cortex | **-0.76 (0.00)** | **1.00** | 0.00 (0.00) | 0.00 |
| TPJ self-connection | 0.00 (0.00) | 0.00 | 0.00 (0.00) | 0.00 |
| auditory cortex to INS | 0.00 (0.00) | 0.00 | 0.00 (0.00) | 0.00 |
| auditory cortex to TPJ | **0.12 (0.00)** | **1.00** | 0.00 (0.00) | 0.00 |
| auditory cortex self-connection | **0.45 (0.00)** | **1.00** | **-0.35 (0.01)** | **1.00** |

Note: Parameters with a posterior probability > 95% were highlighted in bold font. Conditional
covariance for each parameter is given in parentheses.

**Figure 3.**
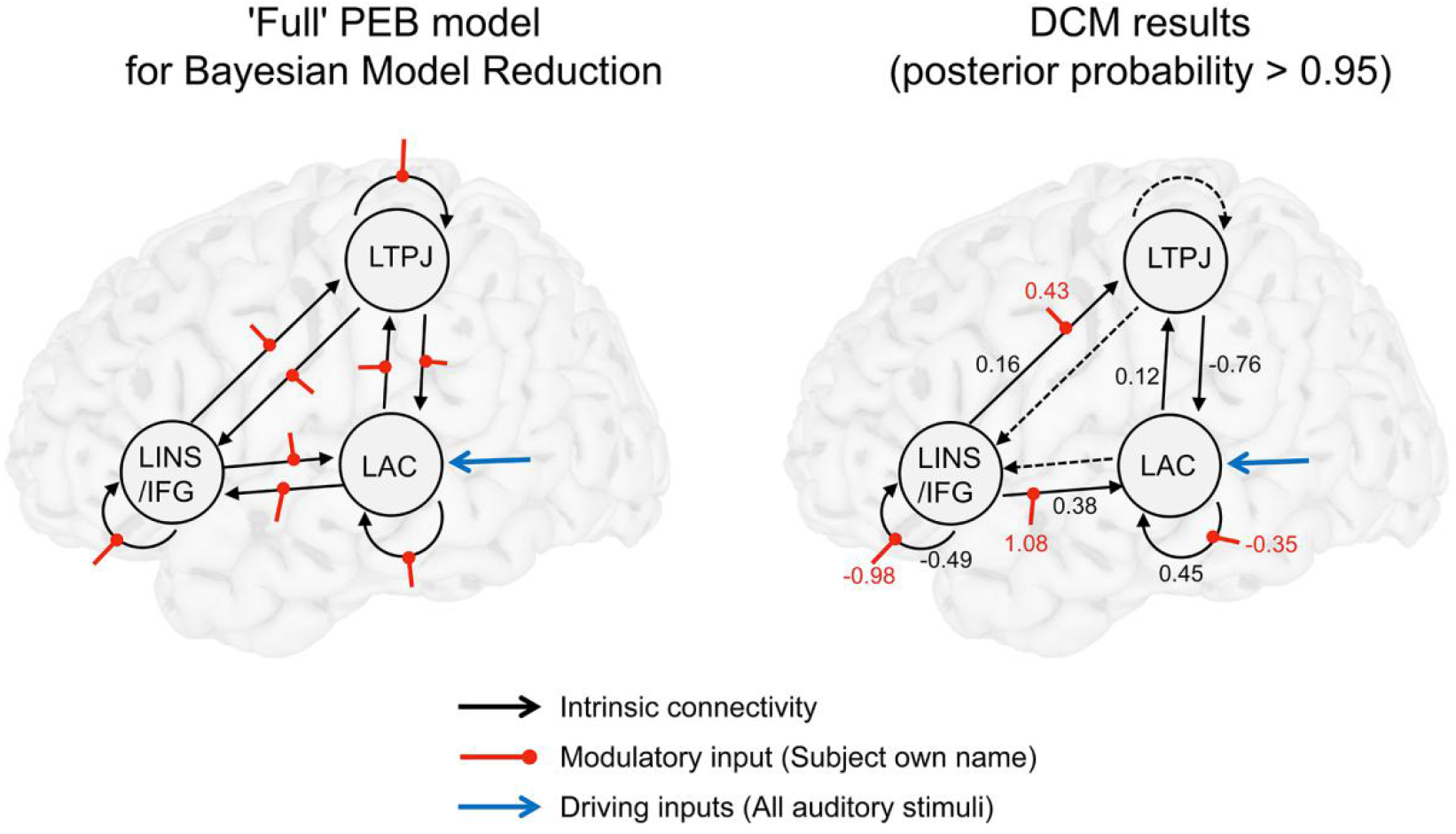
The modulatory effect of SON on effective connectivity. (A) A scheme of the full connectivity model (‘full’ model) for Bayesian model reduction, in which black arrows represent intrinsic connectivity, red dots represent modulatory effect of SON, and blue arrows represent driving inputs using all four experimental stimuli. (B) Group-level DCM results at 95% posterior probability. A solid line represents an intrinsic connectivity with posterior probability over 95%, while a dotted-line represents an intrinsic connectivity with posterior probability below 95%. The red dots represent the modulatory effect of SON with posterior probability over 95%. AC = auditory cortex, INS = insula, IFG = inferior frontal gyrus, TPJ = temporal parietal junction.

## 4. Discussion

Combining the information from activation strength (i.e., GLM) and activation pattern (i.e., MVPA), our study investigated the effective connectivity among multimodal regions (i.e., anterior INS/IFG, TPJ) and unimodal sensory regions (i.e., primary auditory cortex) during SON self-name processing. Here, our GLM results showed that SON consistently induced a higher activation in the left TPJ and bilateral anterior INS/IFG compared with FN, UN, and SONREV, respectively. Using MVPA, we found that the activation pattern of bilateral primary auditory cortex could reliably classify SON and non-SON stimuli. Finally, the DCM results showed that SON increased the effective connectivity from the left anterior INS/IFG to the left primary auditory cortex, and to the left TPJ, as compared with the non-SON conditions, which indicated an excitatory effect of SON (Zhang and Du, 2022). The current findings showed for the first time that SON and non-SON processing could be decoded in unimodal sensory regions (i.e., primary auditory cortex), which cannot be observed with a conventional GLM analysis. Moreover, our results showed that SON processing promoted effective connectivity from the anterior INS/IFG to the primary auditory cortex and TPJ, which highlighted the crucial role of the anterior INS/IFG on SON processing.

Most interestingly, using multivariate analysis, our findings extended prior knowledge by demonstrating a highly significant engagement of the primary auditory cortex during SON processing, which was not observed in previous studies using conventional univariate analysis (Qin et al., 2012; Tacikowski et al., 2011, 2013), including the GLM analysis adopted in the current study. In addition, it was found that the primary auditory cortex also showed a discriminative pattern among each pair of the non-SON stimuli (including FN, UN, SONREV). These findings were supported by previous studies, which found that the activation pattern of the primary auditory cortex was able to classify not only fundamental auditory perception, for instance, multiple sound categories (Staeren et al., 2009; Zhang et al., 2015), but also high-level cognitive processes, including music-induced emotions (Putkinen et al., 2021), perceptual interpretation of ambiguous sound (Kilian-Hütten et al., 2011), and semantic processing (Fedorenko et al., 2012). Taken together, our results provided evidence that bilateral primary auditory cortex contains information to differentiate SON and non-SON, which may contribute to a deeper understanding of the role of the primary auditory cortex in processing a highly self-related stimuli, i.e., SON.

In addition to these differences revealed by activation patterns, our DCM results provide direct evidence of an excitatory effect on the connection from the multimodal anterior INS/IFG to the unimodal primary auditory cortex for the SON condition. As a core region underlying internal sensory integration (Park and Tallon-Baudry, 2014; Seth et al., 2012), the anterior INS/IFG was found to be crucial for self-processing (Critchley and Seth, 2012; Seth, 2013). Evidence from human (Saura et al., 2008) and non-human primates (Hackett et al., 1999) have shown an anatomical connectivity between the left anterior INS/IFG and the auditory cortex. On the other hand, convergent findings have shown that prior knowledge could facilitate speech perception through a top-down excitatory modulation from the left anterior INS/IFG to the lower-level auditory cortex (Di Liberto et al., 2018; Park et al., 2015; Sohoglu et al., 2012). These findings could be explained within the framework of the predictive coding theory, which proposed that perceptual contents could be a result from a hierarchical Bayesian, knowledge-driven inference (Clark, 2013; Friston, 2009; Lee and Mumford, 2003). Our finding supported this theory, and extended the current knowledge by showing a top-down influence from the anterior INS/IFG to the primary auditory cortex modulated by the SON. Considering that self-processing could be a continuously ongoing process that influence one’s experiences in a very fundamental way (Northoff et al., 2006), the current result suggested that our on-going experience of the self could facilitate the recognition of an external self-related stimuli (i.e., SON), when the low-level auditory cortex received an excitatory influence from the high-level anterior INS/IFG.

Furthermore, our DCM results showed an excitatory effect of SON on the connection from the anterior INS/IFG to the TPJ. The left TPJ found in the current study have been widely investigated in previous studies, implicating its crucial role in sensory and sensorimotor integration (Eddy, 2016; Igelström and Graziano, 2017), which was also found to be selectively activated during the self-other distinction task (e.g., mentalizing, or theory of mind) (Knyazev et al., 2021; Tan et al., 2022). Anatomically, the left anterior INS/IFG and TPJ was found to be connected (Saura et al., 2008), and disruption of this connection was observed in patients with personal neglect, in which the individual was unable to perceive the contralesional half of their body (Committeri et al., 2007). Convergent evidence has also indicated that functional integration between the anterior INS/IFG and TPJ was crucial for bodily self-consciousness (Park and Blanke, 2019). Combining the above evidence, the current finding of an excitatory influence from the anterior INS/IFG to TPJ suggested that TPJ might not be involved in the self-other distinction process independently, but requires an excitatory influence from another highly self-related region, i.e., the anterior INS/IFG.

One issue should be noted. Our GLM results failed to show activation in the right TPJ after multiple comparison correction. Furthermore, the DCM results did not show the effective connectivity from the right anterior INS/IFG to the right TPJ modulated by SON. These findings may suggest a lateralized role of the TPJ in self-processing. Considering that it is still debated whether left and right hemisphere was more engaged in self-processing (Qin et al., 2020), which was beyond the scope of this study, further research is needed to answer this question.

## 5. Conclusions

The current study extended our understanding of self-processing in two main aspects. Firstly, we provided evidence that different activation patterns could be reliably detected in bilateral primary auditory cortex between the SON and non-SON stimuli (i.e., FN, UN, and SONREV). Secondly, our DCM results showed that perception of SON strongly enhanced the effective connectivity from the left anterior INS/IFG to the primary auditory cortex, and to the TPJ. Combining previous finding of the involvement of the anterior INS/IFG in interoceptive processing, subjective feelings, and self-processing (Seth, 2013; Seth et al., 2012), the current results suggested that interoceptive processing might facilitate the perception of an external self-related stimuli, by promoting the neural activity in the primary auditory cortex, which offered new insight into the neural mechanism of self-processing.

## Acknowledgements

This work was supported by Key Realm R&D Program of Guangzhou (202007030005), the National Natural Science Foundation of China (31971032), the Major Program of the National Social Science Fund of China (18ZDA293), the Basic and Applied Basic Research Foundation of Guangdong Province (2020A1515011250), Guangdong-Hong Kong-Macao Greater Bay Area Center for Brain Science and BrainInspired Intelligence Fund (2019023), Key Realm R&D Program of Guangdong Province (2019B030335001).

## Author contributions

Conceptualization, Pengmin Qin; Methodology, Hang Wu, and Pengmin Qin; Software, Hang Wu, and Yueyao Liu; Formal Analysis, Hang Wu; Investigation, Dong Wang, and Hang Wu; Resources, Jin Cao, Yujuan Huang, and Mincong Qiu; Writing - Original Draft, Hang Wu; Writing - Review & Editing, Hang Wu, Musi Xie, and Pengmin Qin; Visualization, Hang Wu, Yiwen Wang, and Liwei Zhou; Supervision, Pengmin Qin.

## Data/code availability statement

All data needed to evaluate the conclusions in the paper are present in the paper. Additional data and code related to this paper may be requested from the corresponding author upon reasonable request.

## Supplementary Materials

**Supplementary Table 1.** Parameter Estimates of the Right Hemisphere

| Connection | Intrinsic connectivity | Posterior probability | Modulatory effect of SON | Posterior probability |
| --- | --- | --- | --- | --- |
| INS to TPJ | **0.14 (0.00)** | **1.00** | 0.00 (0.00) | 0.00 |
| INS to auditory cortex | 0.00 (0.00) | 0.00 | **0.58 (0.03)** | **0.99** |
| INS self-connection | **-0.45 (0.00)** | **1.00** | **-0.69 (0.01)** | **1.00** |
| TPJ to INS | **0.11 (0.00)** | **1.00** | 0.00 (0.00) | 0.00 |
| TPJ to auditory cortex | 0.00 (0.00) | 0.00 | 0.16 (0.03) | 0.57 |
| TPJ self-connection | **-0.15 (0.00)** | **0.99** | **-0.78 (0.01)** | **1.00** |
| auditory cortex to INS | 0.00 (0.00) | 0.00 | 0.00 (0.00) | 0.00 |
| auditory cortex to TPJ | 0.00 (0.00) | 0.00 | 0.00 (0.00) | 0.00 |
| auditory cortex self-connection | **0.52 (0.00)** | **1.00** | -0.19 (0.01) | 0.93 |

Note: Parameters with a posterior probability > 95% were highlighted in bold font. Conditional
covariance for each parameter is given in parentheses.

**Supplementary Figure 1.**
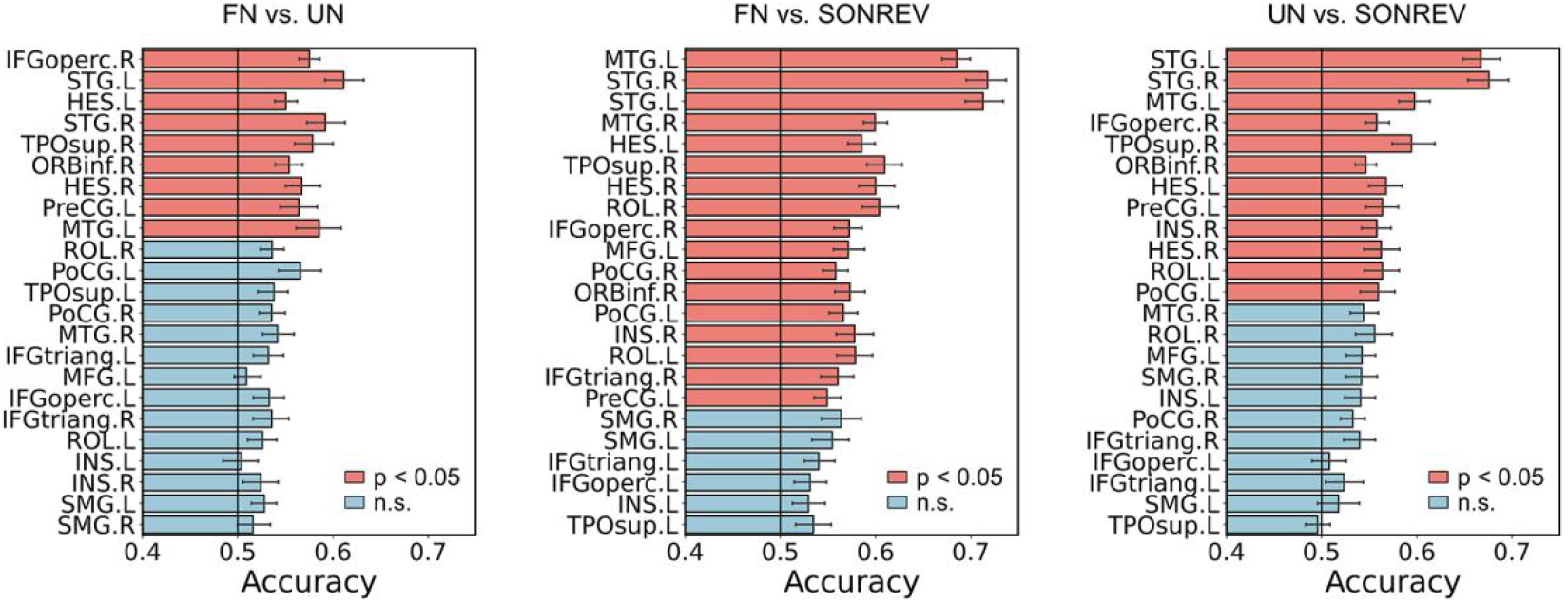
ROI-based MVPA accuracy among the FN, UN and SONREV. Based on the overlapping regions between the searchlight result and AAL 90 atlas, mean accuracy of ROI-level MVPA classification was presented for FN vs. UN, FN vs. SONREV, and UN vs. SONREV, respectively. For each classification, red bars indicate a mean accuracy significantly above chance-level (50%), and blue bars indicate the opposite. SON = subject’s own name, FN = familiar name, UN = unknown name, SONREV = a sound clip with inversed sound waves of the SON.

**Supplementary Figure 2.**
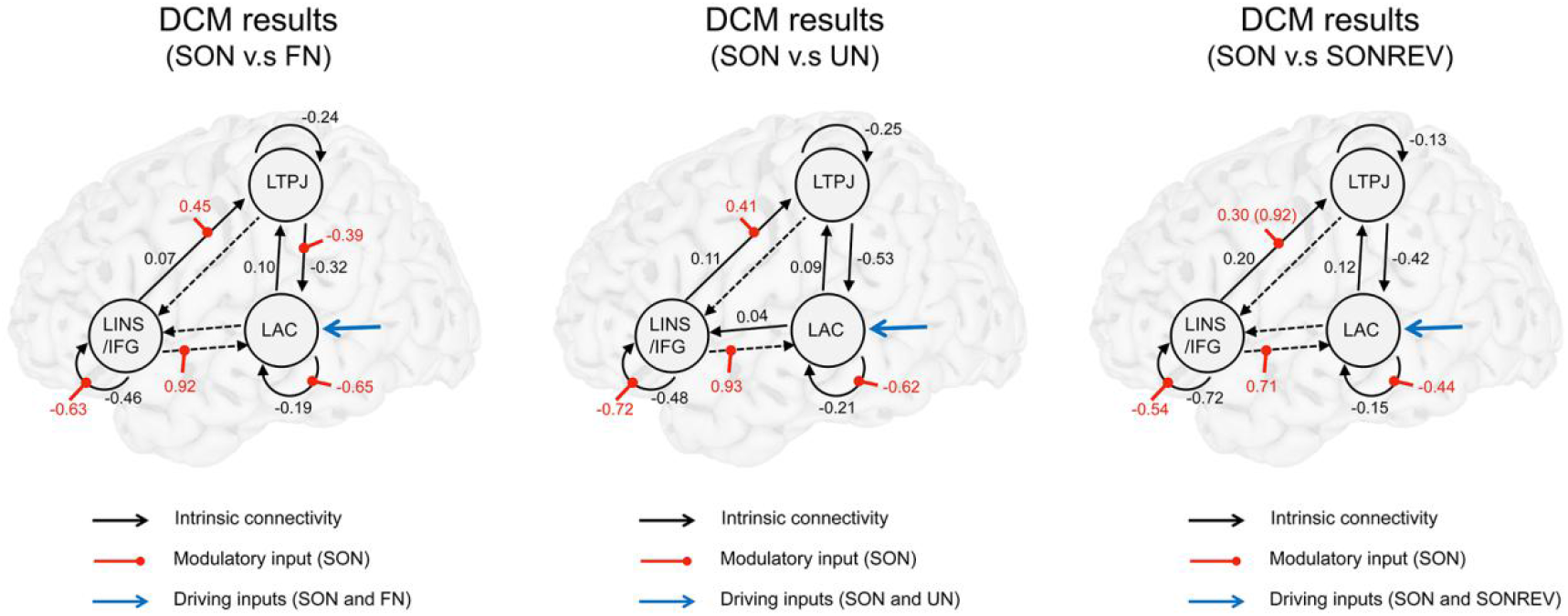
The modulatory effect of SON on effective connectivity relative to the FN, UN, and SONREV, respectively. From left to right, Group-level DCM results were presented for SON vs. FN, SON vs. UN, SON vs. SONREV, respectively, which were controlled at 95% posterior probability. For each DCM model, black arrows represent the intrinsic connectivity, red dots represent the modulatory effect of SON, and blue arrows represent driving inputs. Solid line represents an intrinsic connectivity with a posterior probability over 95%, while dotted-line represents a posterior probability below 95%. AC = auditory cortex, INS = insula, IFG = inferior frontal gyrus, TPJ = temporal parietal junction.

**Supplementary Figure 3.**
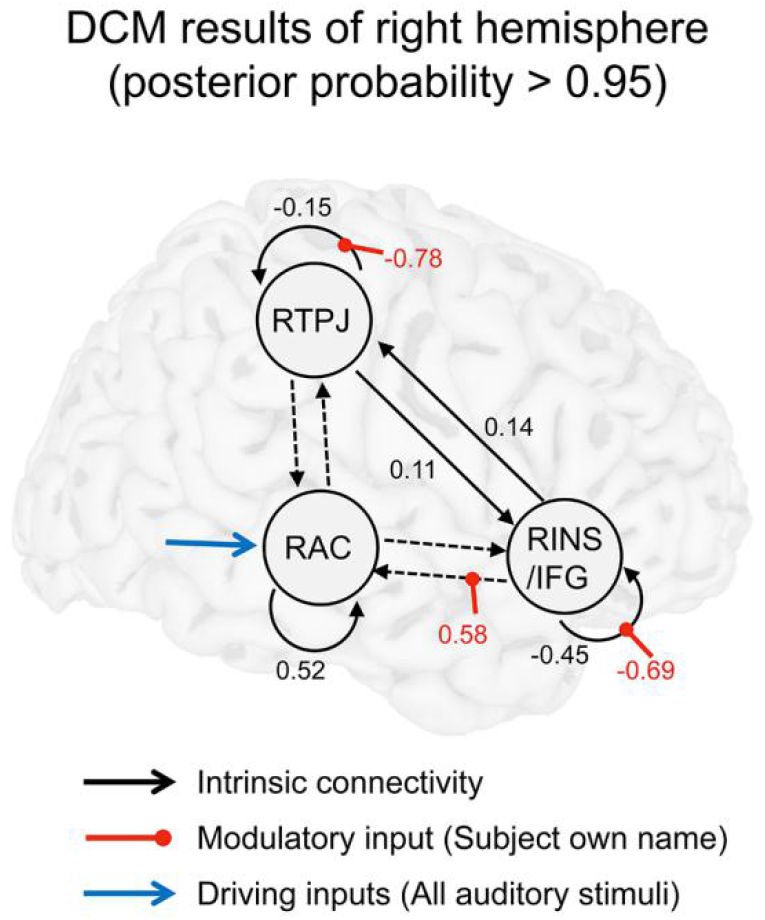
The modulatory effect of SON on the effective connectivity in the right hemisphere. Group-level DCM results was presented at 95% posterior probability, in which black arrows represent the intrinsic connectivity, red dots represent the modulatory effect of SON, and blue arrows represent driving inputs of all four experimental stimuli. Solid line represents an intrinsic connectivity with posterior probability over 95%, while dotted-line represents the opposite. AC = auditory cortex, INS = insula, IFG = inferior frontal gyrus, TPJ = temporal parietal junction.

